# Predicted binding interface between coronavirus nsp3 and nsp4

**DOI:** 10.1101/2022.03.05.483145

**Authors:** Zach Hensel

## Abstract

Double membrane vesicles (DMVs) in coronavirus-infected cells feature pores that span both membranes. DMV pores were observed to have six-fold symmetry and include the nsp3 protein. Co-expression of SARS-CoV nsp3 and nsp4 induces DMV formation, and elements of nsp3 and nsp4 have been identified that are essential for membrane disruption. I describe a predicted luminal binding interface between nsp3 and nsp4 that is membrane-associated, conserved in SARS-CoV-2 during the COVID-19 pandemic and in diverse coronaviruses, and stable in molecular dynamics simulation. Combined with structure predictions for the full-length nsp4 monomer and cryo-EM data, this suggests a DMV pore model in which nsp4 spans both membranes with nsp3 and nsp4 inserted into the same bilayer. This approach may be able to identify additional protein-protein interactions between coronavirus proteins.

## Introduction

Non-structural proteins (nsps) are essential and highly conserved in coronaviruses, yet some lack experimental structural data for nsps and their protein-protein interactions. Structure prediction using AlphaFold2 has recently been shown to accurately predict the structures of human protein-protein interactions (Bryant, Pozzati, and Elofsson 2021), suggesting that this method can fill in gaps in experimental data for viral protein-protein interactions to suggest new experiments, molecular mechanisms, and possible drug targets.

Double membrane vesicles (DMVs) in cells infected by coronavirus were recently found to be a primary site of viral RNA synthesis (Snijder et al. 2020) and contain 6-fold symmetric pores spanning both membranes, consisting in part of nsp3 (Wolff et al. 2020). Co-expression of full-length SARS-CoV nsp3, nsp4, and nsp6 in HEK293T cells induced the formation of DMVs that resemble those in SARS-CoV-infected cells, and membrane rearrangement, but not DMV formation, was observed with an nsp3 C-terminal construct containing putative transmembrane domains (Angelini et al. 2013). It was later shown that SARS and MERS nsp3/4 cleavage is essential for DMV formation, while nsp6 is not needed (Oudshoorn et al. 2017).

Co-localization of nsp3, nsp4, and nsp6, the presence of predicted transmembrane domain regions in each, and the presence of nsp3 in the DMV pore suggest that these could be some of the key components of a functional pore. Association between nsp3 and nsp4 has been mapped to the nsp3 C-terminal region containing predicted transmembrane domains (Hagemeijer et al. 2011) and a short region of nsp4 that can be disrupted with a 2-residue deletion, which also prevents viral replication (Sakai et al. 2017). A model was suggested in which nsp3 and nsp4 insert into opposing membranes and bind in the lumen to bring membranes together, with conserved cysteines playing an important role (Hagemeijer et al. 2014).

This work began with the observation that SARS-CoV-2 Delta variant sequences lacked a 3-residue deletion in nsp6 (Δ106–108) that confers a fitness advantage in several variants of concern (Richard et al. 2021). In place of this deletion, Delta sub-lineages were defined by various nsp6 mutations including L37F that had been identified early in the pandemic (Benvenuto et al. 2020). The most common nsp6 mutations upon the emergence of Delta were V149A and T77A, with sub-lineages subsequently acquiring additional nsp6 mutations. For example, V149A sub-lineages now largely include T181I, and addition of I162V to T77A defined the AY.3 sub-lineage (Hensel 2021; Linsenmeyer et al. 2021). Intriguingly, nsp6 T77 and I162 were adjacent in DeepMind’s prediction of the nsp6 structure made using AlphaFold2 (DeepMind 2020; Jumper et al. 2021), suggesting that recent advances structure prediction could be combined with observing evolution during the pandemic to draw inferences regarding structure and function.

Importantly, AlphaFold2 models are not trained to directly predict the impact of mutations and may not perform well in this task (Pak et al. 2021). However, data on which mutations are possible, which occur together, and which confer fitness advantages or disadvantages can be integrated with structure prediction models trained largely on pre-pandemic data and structural data to begin to answer such questions. The release of the source code and models for AlphaFold2 (Jumper et al. 2021) combined with the fast-paced development of tools such as ColabFold that make this approach accessible to all (Mirdita, Ovchinnikov, and Steinegger 2021).

## Methods

### Structure prediction

Most predictions were generated using AlphaFold2 (Jumper et al. 2021) as implemented in ColabFold (Mirdita, Ovchinnikov, and Steinegger 2021), using the LocalColabFold build available in September 2021. In this implementation, multiple sequence alignments for each monomer in the complex are padded by gap characters and concatenated, as in RoseTTAFold (Baek et al. 2021). Template structures were not used, and predictions were iterated for up to 48 recycles. ColabFold was otherwise used with its default settings including “Turbo” mode and using MMseqs2 for creating multiple sequence alignments (Steinegger and Söding 2017). As implemented on Google Colab, ColabFold is limited to predicting between 1300 and 1400 residues, so monomer predictions, such as those published by DeepMind (DeepMind 2020), were used to guide smaller scale structure predictions that empirically converged in fewer recycle steps in addition to requiring less GPU memory. Preliminary experiments showed that AlphaFold2 models would often converge on similar structures, but without high probability contacts between residues and with low confidence locally at protein-protein interfaces and/or globally, as measured by *plDDT* and *pTMscore*, respectively (Jumper et al. 2021). This subjective observation has since been quantified, with *pDockQ* estimating the quality of a protein-protein interaction prediction from the number of residues in contact and their average *plDDT* (Bryant, Pozzati, and Elofsson 2021). Putative complexes were pursued further if they met three criteria: (1) high *plDDT* at interfaces, (2) high confidence interactions between monomers in distograms, and (3) reproducible in multiple AlphaFold2 models.

Predictions were made for various regions and complexes of nsp3, nsp4, and nsp6. The interaction identified in preliminary experiments that showed consistent results for all models with the highest confidence interactions between subunits was found in predictions combining nsp3 residues 1401–1602 and nsp4 residues 1–123 (**
Fig. 1
**). The recently released AlphaFold-Multimer (Evans et al. 2021) was used as implemented in ColabFold to compare to a differently trained model and found to give similar results for the nsp3-nsp4 complex (**
Fig. 1a
**, lighter color shades), but was more likely to converge on predictions with steric clashes. Both AlphaFold2 and AlphaFold2-Multimer predicted specific interactions between nsp3 and nsp4 residues conserved in SARS-CoV-2 and MHV (**
Fig. 1b
**), with qualitatively similar distograms for all models (**
Fig. 1c
**) and predicted structures that showed strong overlap when aligned to nsp4 for all but the lowest-ranked predictions (**
Fig. 1a
**). Additionally, predictions included a conserved fold for an nsp3 region at the end of this sequence (**
Fig. 1d
**), but without consistently predicted interactions with other nsp3 residues or with nsp4 (**
Fig. 1c
**). Further work is required to determine structures of transmembrane regions of nsp3 and possible protein complexes.

**Figure 1.**
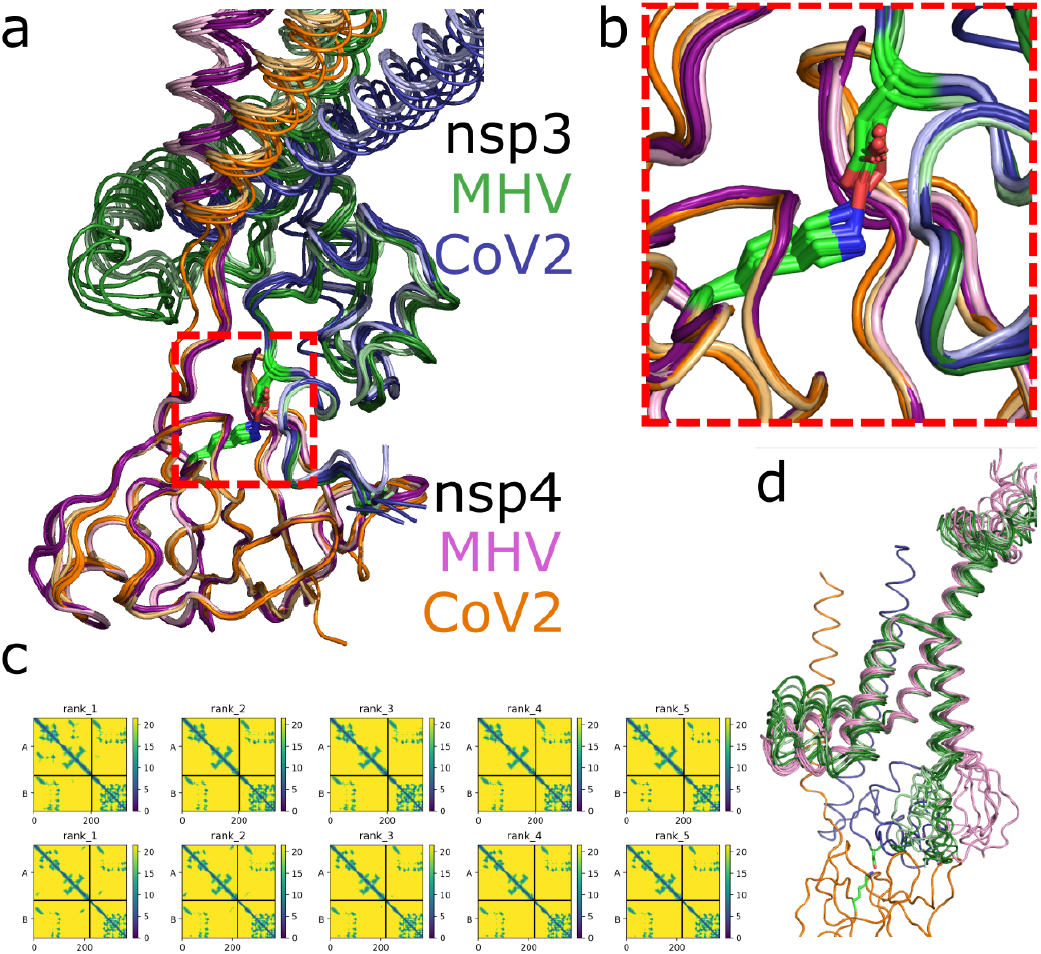
AlphaFold2 and AlphaFold-Multimer predictions for SARS-CoV-2 nsp3 1401–1602 and nsp4 1–123 and equivalent MHV residues. (**a**) SARS-CoV-2 (blue/orange) and MHV (green/magenta) nsp3 and nsp4 predictions are aligned to SARS-CoV2 nsp4 residues 36–123. Both AlphaFold2 (darker shades) and AlphaFold2-Multimer (lighter shades) predictions are shown, giving a total of 18 overlaid predictions (five models each, omitting the lowest scoring SARS-CoV-2 predictions by AlphaFold2 and AlphaFold-Multimer). (**b**) A predicted interaction between an nsp3 aspartate and an nsp4 lysine conserved between SARS-CoV-2 and MHV is highlighted. (**c**) AlphaFold2 distograms (SARS-CoV-2, top; MHV, bottom; predictions ranked by pTMscore) predict contacts between nsp3 and nsp4 transmembrane helices followed by a contacts between nsp3 and residues throughout the first luminal nsp4 domain. Within each distogram, nsp3 appears in the upper left, nsp4 in the lower right, and predicted nsp3-nsp4 contacts in the lower left and upper right. (**d**) SARS-CoV-2 nps3 residues 1491–1602 and its MHV equivalent are aligned to the top scoring SARS-CoV-2 structure and show similar predicted folds in a transmembrane region; nsp3 1401-1490 and nsp4 are only shown for one prediction.

Final predictions were made using LocalColabFold on a Google Cloud Platform instance with an A100 GPU with 48 recycles. Preliminary experiments showed that “Model 3” and “Model 5” in ColabFold consistently converged to the highest confidence predictions (per pTMscore and *plDDT*), with “Model 3” tending to give the highest *plDDT*. These models correspond to training protocols 1.2.1 and 1.2.3 which were trained without templates (Jumper et al. 2021), so having the best performance for template-free prediction is expected. Complexes reported here are on the order of 600 residues and well within the limits of ColabFold (Mirdita, Ovchinnikov, and Steinegger 2021), so each prediction can be reproduced on Google Colab in less than one hour.

In addition to SARS-CoV-2, 10 other coronavirus nsp3-nsp4 complexes were predicted based on a selection previously used to analyze topology predictions for nsp3 (Kanjanahaluethai et al. 2007). Sequences and their UniProtKB identifiers are shown in **
Table 1
**. For nsp3, transmembrane likelihood for full-length nsp3 was predicted by TMHMM 2.0 (Krogh et al. 2001) and used to identify regions containing possible transmembrane sequences and approximately 50 additional flanking residues. From preliminary experiments with SARS-CoV-2 and MHV complexes (**
Fig. 1
**), it was expected that these sequences would only contain small luminal domains, and that there would be no reproducible interactions between nsp3 domains or additional interactions with nsp4, but a wider region was used to possibly identify other possible interactions.

**Table 1.**
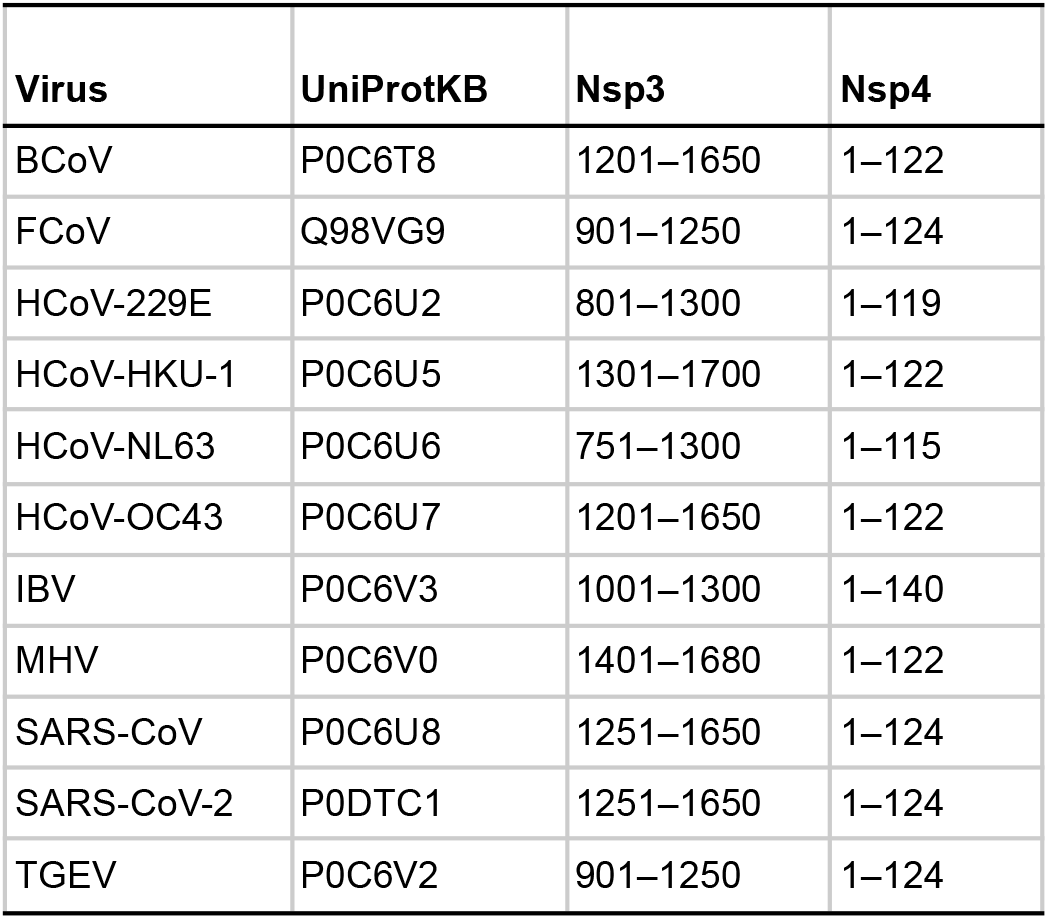
Sequences used for nsp3-nsp4 predictions. For nsp3, sequences were selected by analyzing predicted transmembrane regions in THMM 2.0, including approximately 50 additional surrounding residues. For nsp4, from the first residue to the end of the first predicted luminal domain in a full-length nsp4 prediction, according to a multiple sequence alignment to residue 124 of SARS-CoV-2 nsp4.

### Estimating quality of predicted protein-protein interface

The metric *pDockQ* is an empirical measure used by FoldDock (Bryant, Pozzati, and Elofsson 2021) to estimate the quality of protein-protein interfaces predicted by AlphaFold2. It is defined by a sigmoidal function of four empirical parameters, the number of beta carbons within 8 Å at an interface (*N*), and the average *plDDT* of interface residues (*IF_plDDT*). It is defined by the FoldDock scripts “fetch_plDDT.py” and “vis_analyze.py” as:

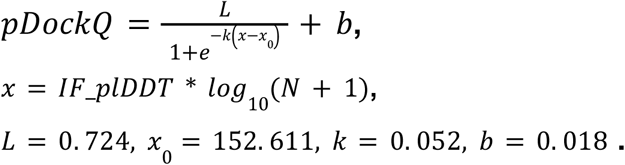

Parameters used for calculating *pDockQ* were retrieved from the FoldDock GitLab repository (9 February 2022), and pDockQ was calculated using a PyMOL script. Following the FoldDock scripts, interface residues were multiply counted both for calculating *IF_plDDT* and *N* if they had multiple interface contacts. The number of unique residues at the interface is not used in calculating *pDockQ*, but is included for reference.

### Molecular dynamics and visualization

Initial molecular dynamics was done using a recently published Google Colab notebook (Arantes et al. 2021) with an OpenMM (Eastman et al. 2017) workflow. Fragments of soluble SARS-CoV-2 nsp3 and nsp4 were simulated by extracting from predicted coordinates nsp3 1428–1488 and nsp4 26–124, truncating sequences N-terminally after predicted transmembrane residues and C-terminally starting where AlphaFold2 had low confidence (low plDDT). Simulation parameters were: ff19SB forcefield (Tian et al. 2020), 150 mM NaCl, NPT ensemble, 310 K, and a 2-fs timestep, with 1000 steps of constrained equilibration followed by a 200 ns of unconstrained molecular dynamics. The simulation contained 37,915 atoms.

To simulate transmembrane nsp3 and nsp4, selected coordinates were expanded to nsp3 1404–1490 and nsp4 1–124. The CHARMM-GUI Membrane Builder (Wu et al. 2014) was used to construct the system with a bilayer consisting of a 48, 20, and 6 POPC, DOPE, and SAPI molecules, respectively, in each leaflet (Olarte et al. 2020). Simulation conditions differing from soluble simulations were: CHARMM36m forcefield (Huang et al. 2017), six rounds of constrained minimization and equilibration, and production runs using hydrogen mass repartitioning with a 4-fs timestep. The simulation consisted of 64,338 atoms and was run for 200 ns using GPU-accelerated OpenMM.

Both simulations included disulfide bonds predicted by AlphaFold2 between all six conserved cysteines in these nsp3 and nsp4 regions. Additionally, it is notable that AlphaFold2 predicts disulfide bridges for all luminal cysteines in full-length nsp4. Figures were arranged and rendered using VMD (Humphrey, Dalke, and Schulten 1996) or Pymol (DeLano 2002). Distograms were adapted from those output by ColabFold or made using NanoHUB (Madhavan et al. 2013). All-atom RMSD was measured following alignment by RMSD minimization to backbone atoms.

### Sequence analysis

Sequences for nsp3 and nsp4 from coronaviruses listed in **
Table 1
** were acquired from UniProt (The UniProt Consortium 2021). Multiple sequence alignments were prepared from full-length nsp3 and nsp4 sequences using Clustal Omega (Sievers et al. 2011), phylogenetic trees were generated from alignments using Simple Phylogeny (ClustalW2), and tree images were generated using iTOL (Letunic and Bork 2021), all with default settings apart from formatting. Sequence logos for nsp3 and nsp4 regions considered in molecular dynamics simulations were generated using WebLogo 3 (Crooks et al. 2004) after acquiring multiple sequence alignments with MMseqs2 following the same workflow used for structure predictions (Steinegger and Söding 2017).

## Results

### A predicted SARS-CoV-2 nsp3-nsp4 interface conserved the pandemic

Based upon preliminary results described above, structures were predicted for heterodimers of SARS-CoV-2 nsp3 1251–1650 and nsp4 1–124 as well as equivalent regions in other coronaviruses (Table 1). An overview of the SARS-CoV-2 nsp3-nsp4 prediction is shown in Fig. 2a. Two protein-protein interfaces were consistently observed to have high confidence predictions (high *plDDT*). As in preliminary predictions for SARS-CoV-2 and MHV, interfaces were observed between transmembrane helices, and between an nsp3 loop that interacts with several regions of the first predicted nsp4 luminal domain (**
Fig. 2a
**), with membrane topology suggested by surface charge and hydrophobicity (**
Fig. 2b
**).

**Figure 2.**
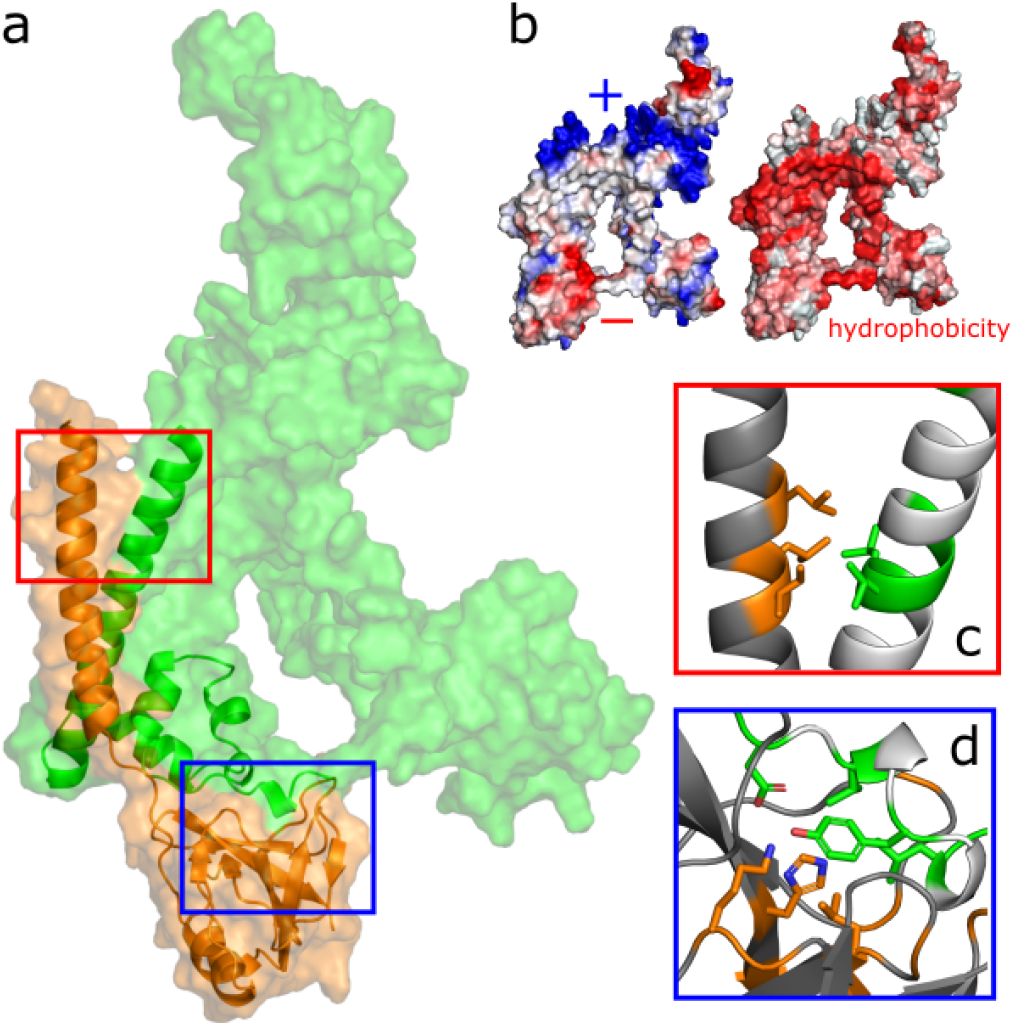
Predicted SARS-CoV-2 nsp3-nsp4 binding interfaces. (**a**) A transparent surface representation of all predicted nsp3 (green) and nsp4 (orange) residues is shown, with cartoon representations indicating regions of nsp3 and nsp4 predicted to interact. (**b**) Vacuum electrostatic potential and sidechain hydrophobicity are shown for the predicted nsp3-nsp4 heterodimer. (**c, d**) Conservation of interacting residues. All residues mutated in fewer than 200 sequences in the CoV-GLUE database were highlighted, and sidechains are shown that also have an atom within 3 Å of any atom in the binding partner. Residues with more than 200 mutated sequences in CoV-GLUE are colored white and gray for nsp3 and nsp4, respectively. (**c**) In the predicted transmembrane interaction, conserved nsp3 (L1417 and L1418) and nsp4 (L7, L10, and I11) residues are adjacen. (**d**) Residues nsp3 D1478 and nsp4 K67 have charged side chains separated by 2.8 Å. Residues nsp3 L1480, Y1483, and L1486 form a hydrophobic interface with nsp4 L90 and H120.

One expects an accurate structure prediction for an essential protein-protein interface to show conservation in sequences. However, this approach is difficult to use for evaluating AlphaFold2 predictions since sequence alignments are used to generate predictions. Millions of viral genome sequences collected during the COVID-19 pandemic provide an alternative approach based upon the relative degree of conservation in sequences not used to make predictions. The CoV-GLUE database (Singer et al. 2020) was queried to find the frequency of all non-synonymous SNPs for nsp3 and nsp4 in the first ~3M SARS-CoV-2 genomes in the database. A cut-off of less than 200 sequences exhibiting any non-synonymous was empirically used to highlight conserved residues, finding that interfaces contained conserved, hydrophobic residues, shown in **
Fig. 2c
** and **
Fig. 2d
**. This interface includes nsp4 H120, which was identified to be part of an essential region for interaction with nsp3 (Sakai et al. 2017).

In **
Table 2
**, nsp3-nsp4 complex predictions are additionally evaluated by *pDockQ* (Bryant, Pozzati, and Elofsson 2021) and estimated binding affinity using the PRODIGY server (Xue et al. 2016). The first method estimates protein-protein interface quality based upon the number and local confidence (*plDDT*) of interface residues. The second has a small contribution from the composition of non-interface residues, with estimated affinities largely determined by the types of contacts made by charged and polar residues. For SARS-CoV-2 nsp3-nsp4, *pDockQ* is ~0.7 using two different AlphaFold2 parameter sets, indicating a prediction of medium quality (Basu and Wallner 2016).

**Table 2.**
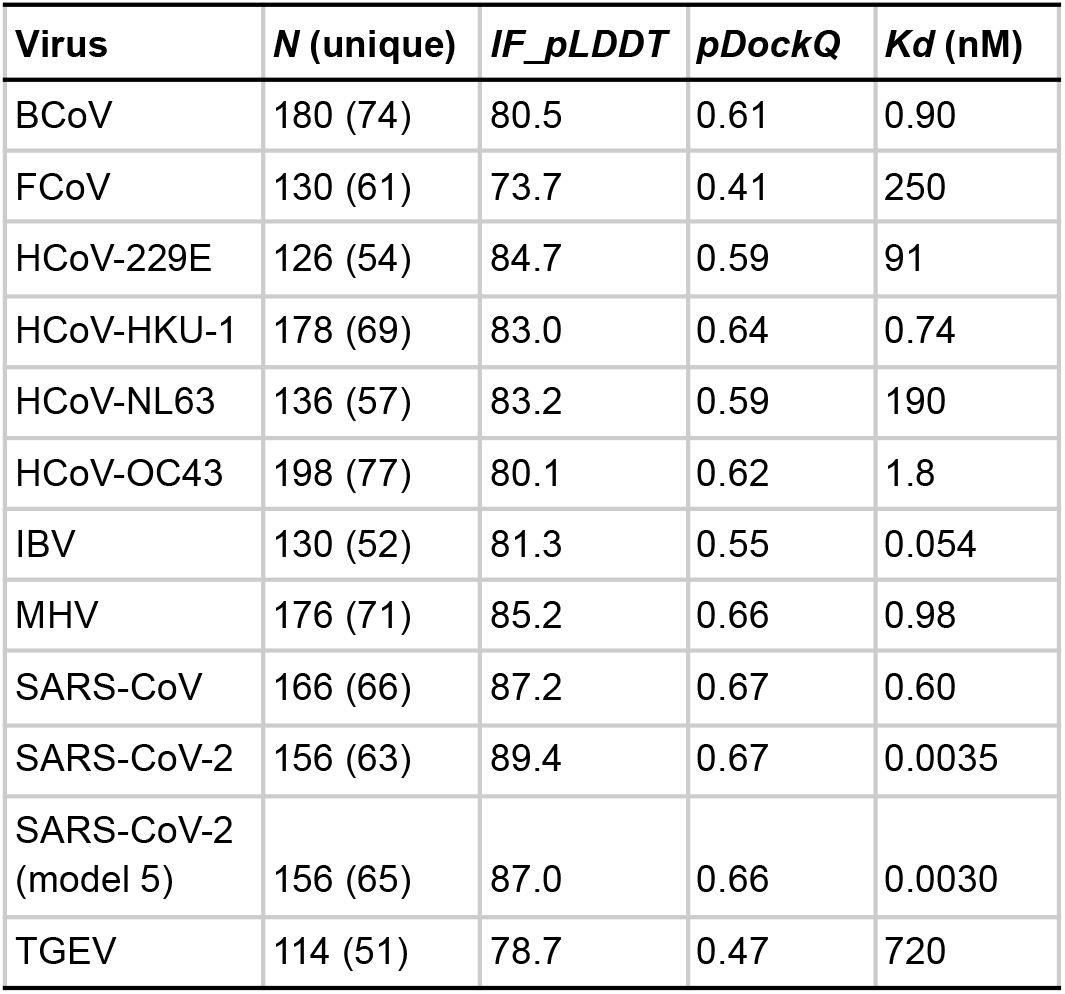
Sequences used for nsp3-nsp4 predictions. For nsp3, sequences were selected by analyzing predicted transmembrane regions in THMM 2.0, including approximately 50 additional surrounding residues. For nsp4, from the first residue to the end of the first predicted luminal domain in a full-length nsp4 prediction, according to a multiple sequence alignment to residue 124 of SARS-CoV-2 nsp4.

### Conserved nsp3-nsp4 interaction predicted in coronaviruses

Based upon preliminary observations of similar predictions for MHV and SARS-CoV-2, predictions were made for diverse coronaviruses to see how widely conserved the predicted nsp3-nsp4 interaction is (**
Table 1
**). All predictions gave *pDockQ* values predicting structures of acceptable or medium quality (**
Table 2
**).

For all complexes, a conserved predicted fold was observed for the first nsp4 luminal domain. Local confidence (*plDDT*) was high for this domain. **
Figure 3
** shows isolated predictions for nsp4 from its first residue to the end of the first luminal domain. Little variation is observed in prediction other than flexibility in the relative orientation of the transmembrane helix and luminal domain.

**Figure 3.**
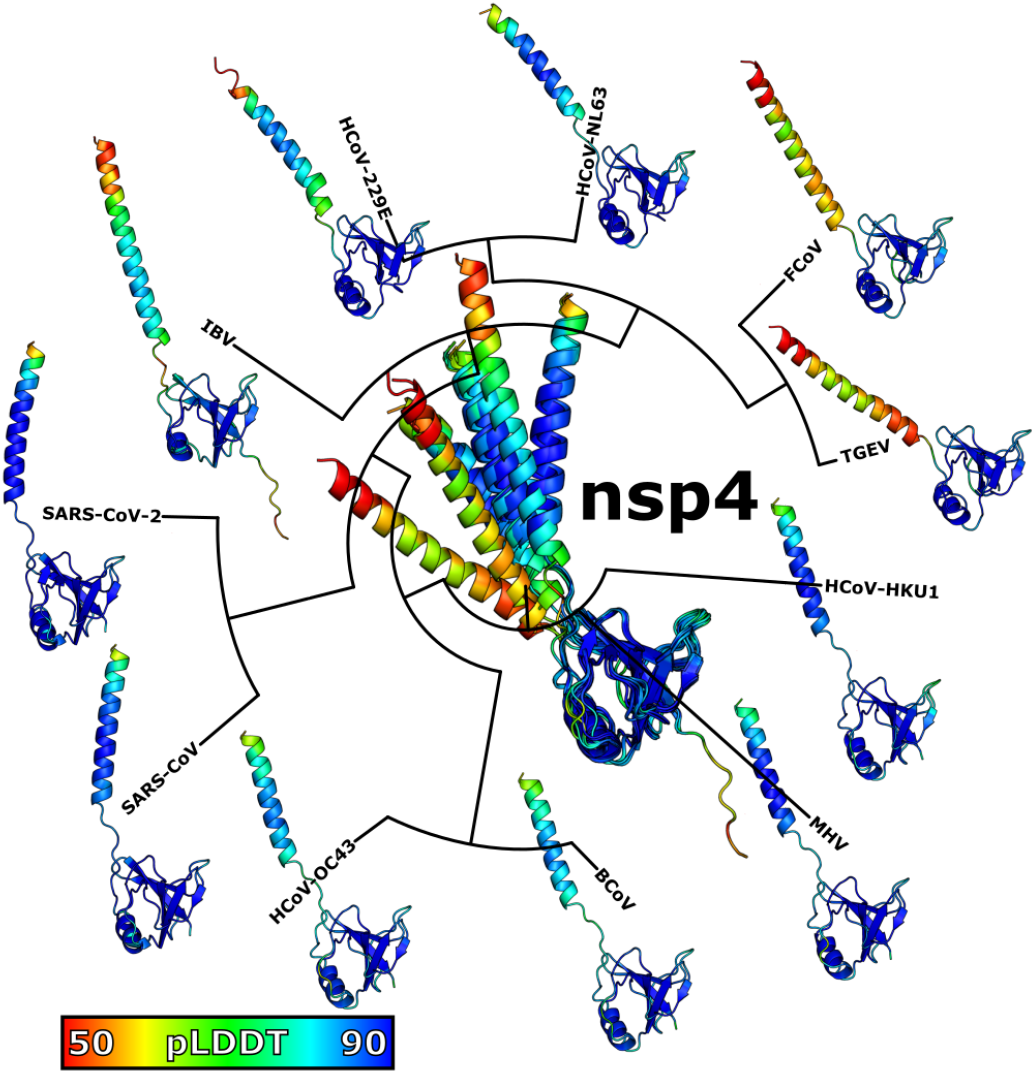
Conservation of predicted nsp4 structure. Only nsp4 is shown from predictions of nsp3-nsp4 complexes, colored by *plDDT*. All predictions are aligned to the SARS-CoV-2 nsp4 luminal domain. Individual predictions are shown against a phylogenetic tree calculated from full-length nsp4 sequences.

Conversely, nsp3 predictions showed wider variation, but shared a region consisting of a transmembrane helix and a small, helical luminal domain ending in a loop predicted to interact with nsp4. In **
Figure 4
**, all predictions are aligned to the nsp4 luminal domain, and nsp3 structures are compared in the region interacting with nsp3. The loop interacting with nsp4 is both the most similar region and that with the highest *plDDT* across all predictions (see inset in **
Fig. 4
** overlaying all predictions). Variation of other parts of nsp3 is observed along the phylogenetic tree, most clearly comparing alphacoronavirus TGEV to betacoronavirus HKU1.

**Figure 4.**
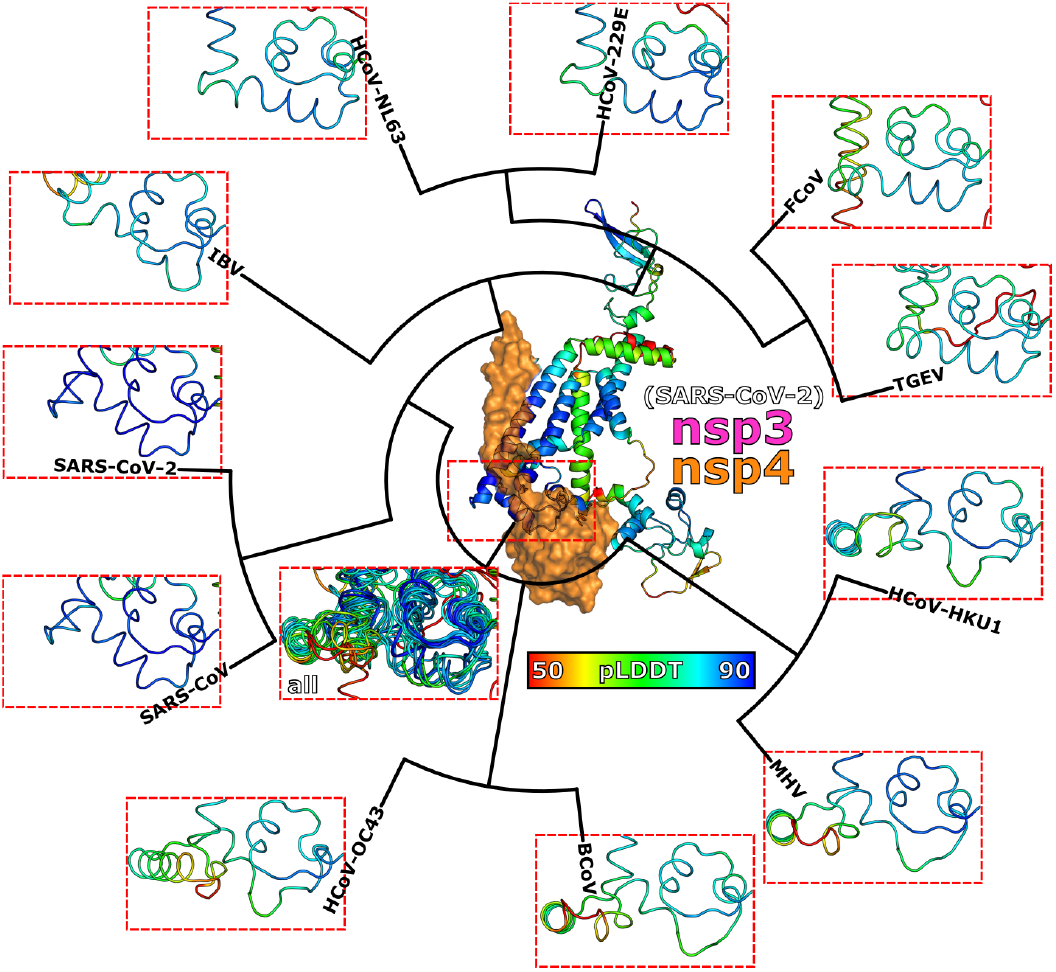
Conservation of nsp3 structure. The center image shows the SARS-CoV-2 nsp3-nsp4 prediction with nsp4 in a surface representation and nsp3 colored by *plDDT*. The region of nsp3 within the red dashed line is shown for each prediction against a phylogenetic tree calculated from full-length nsp3 sequences. The additional image shows all nsp3 predictions overlaid after alignment to the nsp4 luminal domain.

### Molecular simulation of SARS-CoV-2 nsp3 and nsp4 shows key interactions

High-confidence protein-protein complex structures do not necessarily indicate that this structure is stable in the absence of additional components. A series of molecular dynamics simulations were conducted to test whether protein-protein interfaces were maintained and included the same sidechain contacts after equilibration. First, all-atom molecular dynamics was performed for only 10 ns of an nsp3-nsp4 membrane complex (**
FIg. 5a
**) in order to determine likely soluble domains. Second, a 200-ns simulation was performed for predicted soluble regions (nsp3 1428–1488 and nsp4 26–124). This simulation was observed to approach an equilibrium all-atom RMSD of ~2.2 Å within 100 ns (**
Fig. 5b
**) and resulted in a contact map after 200 ns that is qualitatively similar to that predicted by AlphaFold2 (**
Fig. 5c
**). Combined with EM data for a DMV pore (Wolff et al. 2020) and a full-length nsp4 prediction, this suggests a model in which nsp4 spans the luminal space between DMV bilayers (**
Fig. 5d
**).

**Figure 5.**
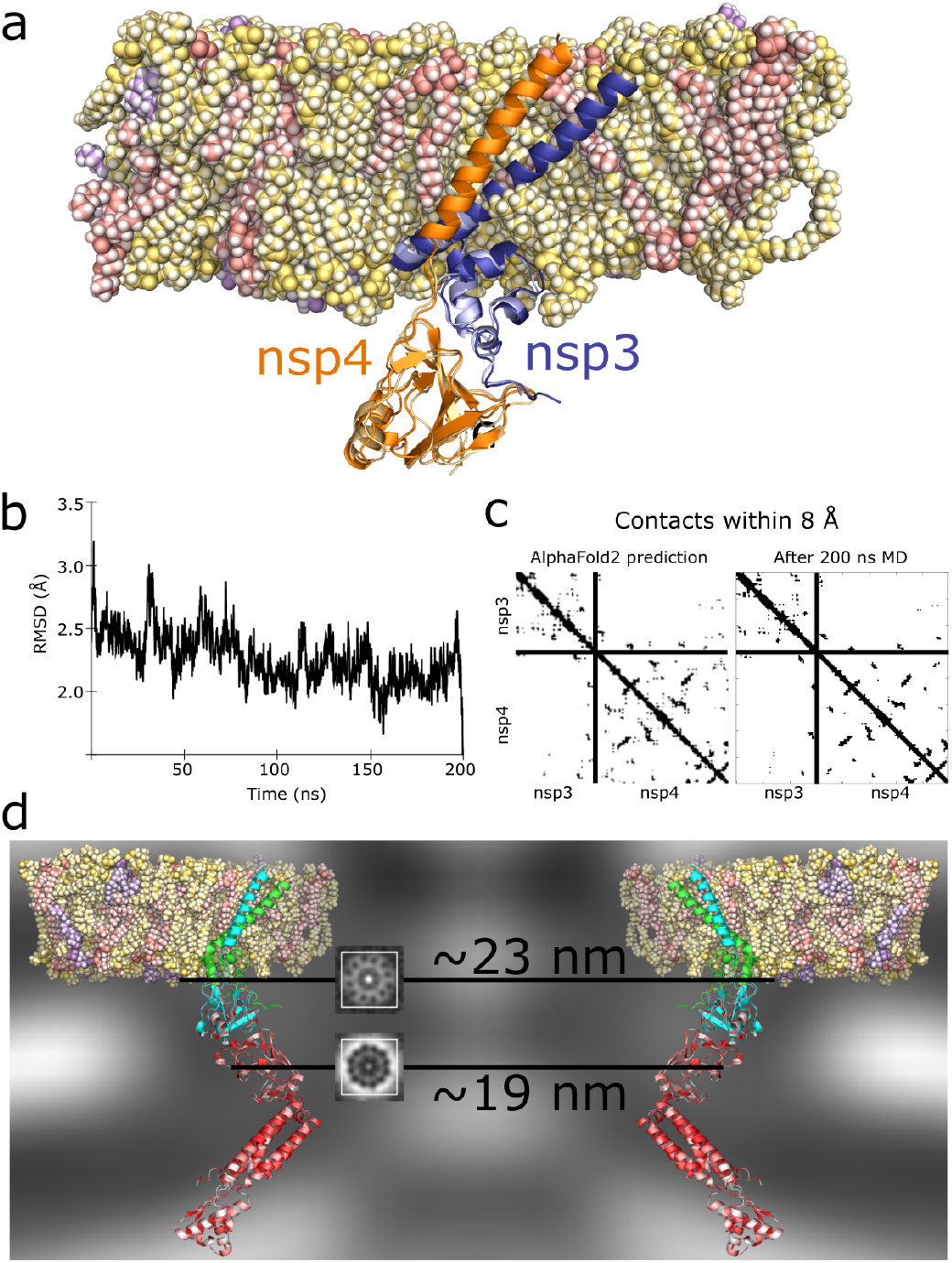
Molecular dynamics results. (**a**) Final frame of 10-ns simulation of nsp3 1404–1490 (blue) and nsp4 1–124 (orange) in a lipid bilayer of POPC (yellow), DOPE (pink), and SAPI (purple); foreground lipids are omitted and the final frame of a 200-ns simulation of nsp3 1428–1488 and nsp4 26–124 (lighter colors) is aligned to the transmembrane structure. (**b**) All-atom RMSD compared to the final frame for the 200-ns simulation. (**c**) The contact map for nsp3-nsp4 from the AlphaFold2 prediction (showing only nsp3 1428–1488 and nsp4 26–124) compared to the contact map after simulating for 200 ns. (**d**) Structure prediction results compared to Cryo-EM of the DMV pore of MHV (EMD-11514; cross-section in background and insets showing slices that include a 12-fold symmetric ring). A full-length nsp4 AlphaFold2 prediction is aligned to nsp4 1–124 and colored by side chain hydrophobicity.

A longer simulation of the membrane complex was conducted to determine which conserved residues in nsp3 and nsp4 define the interface and whether the interface is stable when associated with the membrane. **Movie S1** includes two views of this 200-ns simulation. In **
Fig. 6a
**, sequence conservation for nsp3 1404–1490 is shown along with the average RMSD for these residues across 50 ns of simulation time (after 100 ns of equilibration and before the final 200-ns time point used for alignment). When aligned to the nsp4 transmembrane helix, minima are observed for nsp3 at residues 1424 and 1476, corresponding to the center of the transmembrane helix and a short helix preceding the loop interacting with nsp4. When aligned to nsp4’s luminal domain, a deep minima is observed for a conserved region starting with residue D1478. **
Fig. 6b
** shows sequence conservation for nsp4, and **
Fig. 6c
** shows hydrophobic interfaces formed between nsp3 and nsp4 for residues indicated in **
Fig. 6a
** and **
Fig. 6b
**. Although it is not one of the residues closest in contact with nsp3, nsp4 H120 remains at or near the interface throughout the simulation. Conversely, nsp4 F121 is not at the interface, but is part of the hydrophobic core of nsp4. It is unclear whether the observed effect of nsp4 120–121 deletion (Sakai et al. 2017) comes from eliminating one or both of these features.

**Figure 6.**
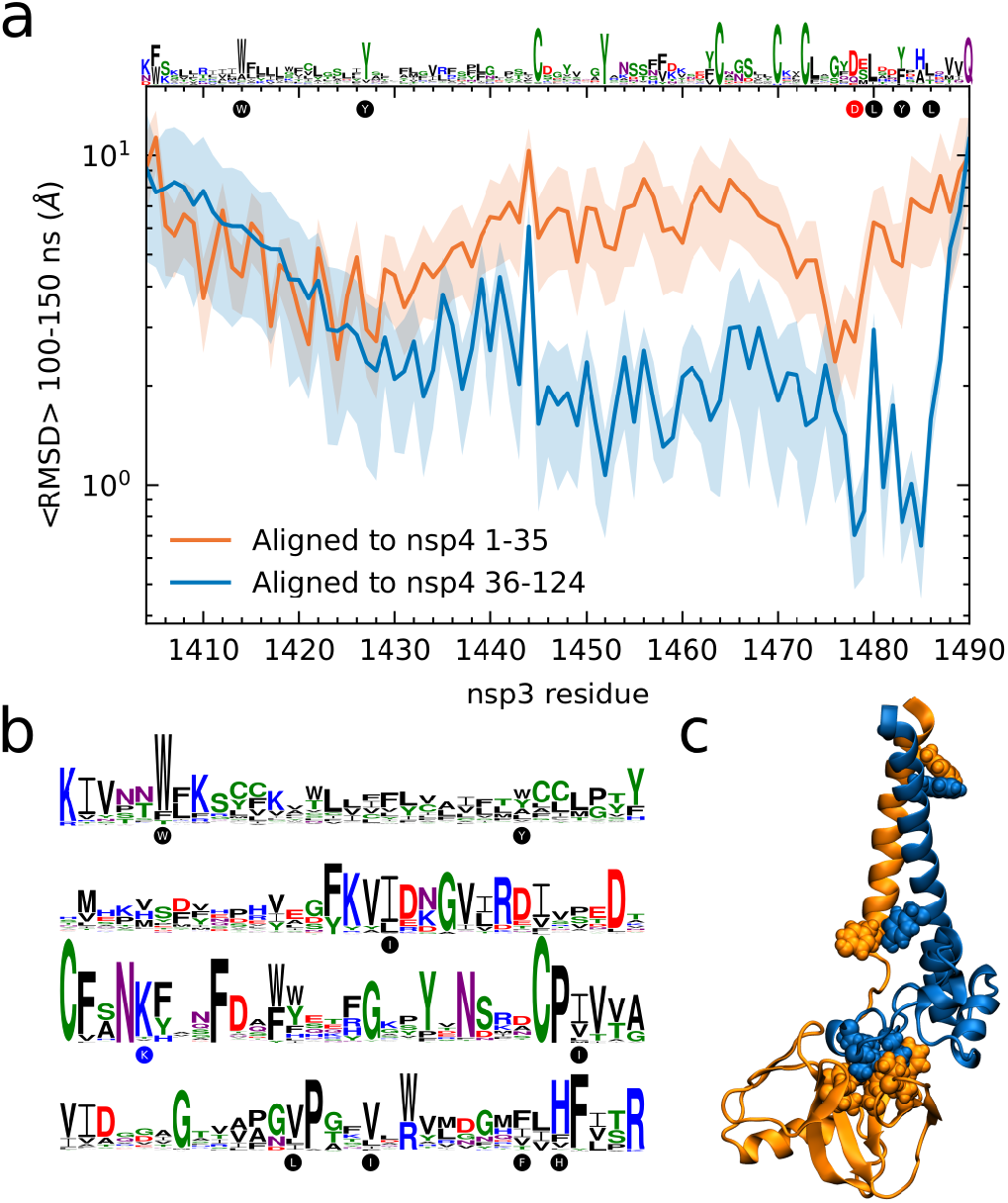
A conserved hydrophobic interface between nsp3 and nsp4. (**a**) Top, a sequence logo for multiple alignment of SARS-CoV-2 nsp3 1404–1490. Circles show SARS-CoV-2 identities for residues highlighted in **c** and discussed below. Bottom, average backbone RMSD (lines) and its standard deviation (shaded) per residue for nsp3 for timepoints between 100 and 150 ns, after alignment to either nsp4 transmembrane (1–35) or luminal (36-124) residue. (**b**) Sequence logo for multiple alignment of SARS-CoV-2 nsp4 1–124. (**c**) Snapshot of nsp3 (blue) and nsp4 (orange) at 100 ns with spheres showing alpha carbon and sidechain atoms for conserved hydrophobic residues highlighted in sequence logos.

The clearest interaction other than hydrophobic interactions is the salt bridge between nsp3 D1478 and nsp4 K67 that was predicted for both SARS-CoV-2 and MHV in initial tests (**
Fig. 1b
**). Charged groups remain separated by less than 3 Å throughout the simulation (**
Fig. 7
**) following fluctuations in the first 30 ns. Interestingly, while nsp4 K67 is highly conserved (**
Fig. 6b
**), nsp3 D1478 is less well conserved (**
Fig. 6a
**). How this interaction varies in other coronaviruses lacking an equivalent to D1478 remains an open question.

**Figure 7.**
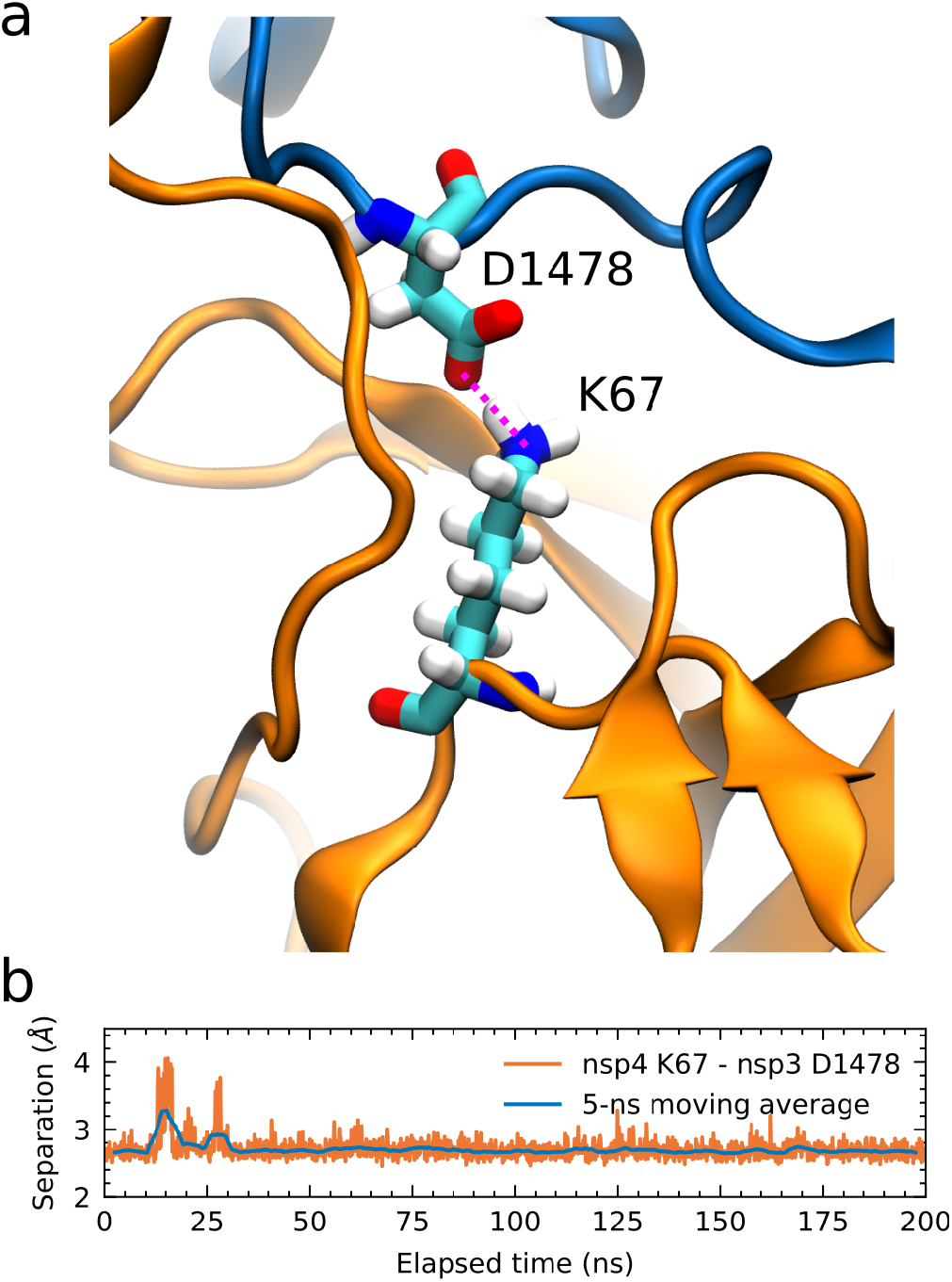
A stable salt bridge between nsp3 and nsp4. (**a**) A snapshot after 100 ns of simulation with a salt bridge between nsp3 D1478 and nsp4 K67 highlighted. (**b**) After aligning the trajectory to nsp4 36–124 at 200 ns, separation between the oxygen and nitrogen atoms highlighted above was calculated along with a 5-ns moving average.

## Discussion

These predictions demonstrate a conserved predicted interface between nsp3 and nsp4. Despite many attempts, no predicted interactions were found between nsp3 and other regions of nsp4 or between the first nsp4 transmembrane helix and its larger predicted transmembrane domain. This suggests a structure similar to that shown in **
Fig. 5d
**, in which nsp4 interacts with nsp3, presumably in the outer DMV bilayer, and spans the luminal space with its large transmembrane domain in the inner DMV bilayer. Speculatively, the 12-fold symmetric density observed in DMV pores for MHV (Wolff et al. 2020) could arise from luminal nsp4 domains.

Determining the structure of complexes including nsp4 may shed light on selection of nsp4 mutants in the COVID-19 pandemic. For example, nsp4 T492I is a mutation found in significant variants. It was most commonly found in B.1.1.519 and B.1.575 in North America in winter 2020/21. It then appeared in an increasing fraction of B.1.617.2 (Delta) sublineages as well as in B.1.621 and C.37 that were spreading rapidly in the first half of 2021 in South America (Mullen et al. 2021). Most recently, it is a defining mutation in Omicron variant lineages. Despite its apparent importance, the functional role of this mutation is unknown.

A limitation of this approach is that it does not predict post-translational modifications. For example, it was observed that mutation of glycosylated nsp4 residues induced defects in viral replication (Gadlage et al. 2010), and this may indicate protein-protein interactions or monomer structures that cannot be accurately predicted in the absence of accounting for post-translational modifications. An additional limitation is that molecular dynamics simulations here are still relatively short, and it is likely that these nsp3 and/or nsp4 regions would have additional protein-protein interactions in the context of a DMV pore. While some interactions are predicted for additional transmembrane nsp3 regions, it was not possible to construct a larger complex with any confidence. Perhaps additional sequence sequence information acquired during the pandemic can be used to improve predictions for SARS-CoV-2 structures and protein-protein interactions.

Lastly, identification of a short nsp3 peptide that may be essential for DMV pore formation suggests a possible therapeutic approach. However, a drug mimicking this region of nsp3 would have to overcome the apparent strength of the interface (**
Table 2
**), ER permeability, and likelihood of evolving resistance.

## Acknowledgments

ZH received support for this work from Project LISBOA-01-0145-FEDER-007660 (Microbiologia Molecular, Estrutural e Celular) funded by FEDER funds through COMPETE2020–Programa Operacional Competitividade e Internacionalização (POCI) and by national funds through Fundação para a Ciência e a Tecnologia (FCT), from a joint research agreement with the Okinawa Institute of Science and Technology, and from the Google Cloud Research Credits program with the award GCP20210916.

This work depends on many open source projects that were indirectly utilized via software cited below. SARS-CoV-2 genome data was not directly accessed in this work, but this analysis would be impossible without rapid sequence publication to GISAID (Bogner et al. 2006). Particularly, I thank Alan Tsang and colleagues in Hong Kong, Sikhulile Moyo and colleagues in Botswana, and Tulio de Oliveira and colleagues in South Africa for rapidly depositing sequencing data for the Omicron variant, and Dakota Howard and colleagues for depositing the first AY.3 sequences.

